# Optimization of Whole Mount RNA multiplexed in situ Hybridization Chain Reaction with Immunohistochemistry, Clearing and Imaging to visualize octopus neurogenesis

**DOI:** 10.1101/2022.02.24.481749

**Authors:** Ali M Elagoz, Ruth Styfhals, Sofia Maccuro, Luca Masin, Lieve Moons, Eve Seuntjens

## Abstract

Gene expression analysis has been instrumental to understand the function of key factors during embryonic development of many species. Marker analysis is also used as a tool to investigate organ functioning and disease progression. As these processes happen in three dimensions, the development of technologies that enable detection of gene expression in the whole organ or embryo is essential. Here, we describe an optimized protocol of whole mount multiplexed RNA *in situ* hybridization chain reaction version 3.0 (HCR v3.0) in combination with immunohistochemistry (IHC), followed by fructose-glycerol clearing and light sheet fluorescence microscopy (LSFM) imaging on whole-mount *Octopus vulgaris* embryos. We developed a code to automate probe design which can be applied for designing HCR v3.0 type probe pairs for fluorescent *in situ* mRNA visualization. As proof of concept, neuronal (*Ov-elav*) and glial (*Ov-apolpp*) markers were used for multiplexed HCR v3.0. Neural progenitor (*Ov-ascl1*) and precursor (*Ov-neuroD)* markers were combined with an immunostaining for phosphorylated-histone H3, a marker for mitosis. After comparing several tissue clearing methods, fructose-glycerol clearing was found optimal in preserving the fluorescent signal of HCR v3.0. The expression that was observed in whole-mount octopus embryos matched with the previous expression data gathered from paraffin-embedded transverse sections. Three-dimensional reconstruction revealed additional spatial organization that had not been discovered using two-dimensional methods.

## 1 Introduction

The recently increased availability of genomic information has spurred molecular research on several cephalopod species, including *Octopus vulgaris* (Albertin *et al*., 2015; Kim *et al*., 2018; Zarrella *et al*., 2019; Li *et al*., 2020). *Octopus vulgaris* or the common octopus, is a cosmopolitan species, and has been the subject of many seminal studies of neural anatomy and behavior (Young, 1971, 1983; Fiorito, Von Planta and Scotto, 1990; Amodio and Fiorito, 2013). How the octopus has expanded its brain and how the nervous system is able to generate these complex cognitive behaviors is a matter of growing research interest.

As novel features arise during the development of organisms, studying embryonic development of the nervous system in cephalopods can give important insights into these research questions. *Octopus vulgaris* spawns several hundreds of thousands of small-sized eggs that develop, depending on the water temperature, in roughly 40 days to independently feeding and swimming paralarvae (Naef, 1928; Deryckere *et al*., 2020). During this period, the central brain develops from placodes to cords and lobes, which represent the adult brain lobes, although only containing about 200,000 cells. Despite the huge difference in size, this larval brain is able to control a number of innate behaviors. Our recent work showed that the larval brain derives from a neurogenic zone located around the eye placode that expresses transcription factors typical for neurogenesis across species (Deryckere *et al*., 2021).

The study of spatial gene expression has been instrumental in defining the molecular patterning and gene function during embryogenesis. In non-model species such as cephalopods, antibody tools are not readily available and often too expensive to develop. Methods that allow detection of mRNA expression *in situ* are more widely applicable. The recent development of *in situ* hybridization chain reaction version 3.0 (HCR v3.0) offers a robust, sensitive, versatile and low-cost method for simultaneous detection of multiple mRNAs in cells or tissues of any organism (Choi, Beck and Pierce, 2014; Choi *et al*., 2016, 2018; Schwarzkopf *et al*., 2021). The method seems to outcompete traditional colorimetric *in situ* hybridization because of its robustness and the option for multiplexing, and other branched DNA probe methods such as RNAscope because of the much lower cost, despite the latter being highly sensitive and easier in use (Jones and Howat, 2020).

In order to follow up organ morphogenesis, technologies have been developed that allow three-dimensional (3D) imaging of whole embryos or organs, often combined with marker gene labeling techniques. Besides classical confocal microscopy, that allows for high resolution imaging at the cellular level, light sheet fluorescence microscopy (LSFM) revolutionized imaging speed of optically transparent organisms including several aquatic embryonic and larval specimen (Santi, 2011). In addition, several methods have been developed to optically clear fixed tissue samples using organic solvent-based methods (e.g. iDISCO+, uDISCO, and BABB) or water-based methods (e.g. CUBIC, Fructose-Glycerol, and TDE) (Richardson and Lichtman, 2015). While these methods often preserve the fluorescent signals generated by genetic labeling in transgenic animal lines, or after immunohistochemistry, currently there is no publication presenting the compatibility of these clearing methods with HCR v3.0 treated cephalopod samples.

Organ development, such as the nervous system, can be complex to understand only using two-dimensional (2D) imaging. 3D imaging can provide an additional perspective. Here, we add to the existing methodology an automation of probe design, and an optimized clearing protocol that retains the signal generated by HCR v3.0 in whole mount *Octopus vulgaris* embryos, even in combination with immunohistochemistry. These methods will advance gene expression analysis in non-model species such as cephalopods. Moreover, we present data on developmental stage XV as experimental stage, which is mid-organogenesis, and shows eye pigmentation that needs to be removed in order to visualize the brain by LSFM. We also include immunohistochemistry to prove that sequential detection of mRNA and protein is feasible using our combined method. Our data confirmed previous findings using HCR v3.0 on transverse sections (Deryckere *et al*., 2021; Styfhals *et al*., 2022) and showed the power of this technique to map phases of organogenesis in 3D.

## 2 Materials and Methods

### 2.1 Animals

Live *Octopus vulgaris* embryos were received from the Instituto Español de Oceanografía (IEO, Tenerife, Spain). Embryos were incubated until reaching the developmental stage XV in the closed standalone system located at the Laboratory of Developmental Neurobiology (KU Leuven, Belgium). The size of an octopus egg, from the stalk till micropyle, is 2 mm x 0,7 mm and a stage XV octopus embryo, from the top of the mantle till the end of arms, is approximately 1,25 mm x 0,88 mm. The stage XV embryos were fixed in 4% paraformaldehyde (PFA) in phosphate buffered saline (PBS) overnight, followed by a wash of Diethyl pyrocarbonate-treated phosphate buffered saline (PBS-DEPC). Embryos were manually dechorionated using tweezers (Dumont #5 Forceps - Biology/Inox, FST) in PBS-Tween (PBST). Embryos were dehydrated into 100% Methanol (MeOH) following a series of graded MeOH/PBST washes, each for 10 minutes: 25% MeOH / 75% PBST, 50% MeOH / 50% PBST, 75% MeOH / 25% PBST, 100% MeOH, 100% MeOH. Dehydrated embryos were kept at -20 °C overnight or until further use.

### 2.2 *In situ* Hybridization Chain Reaction version 3.0 with(out) Immunohistochemistry

#### 2.2.1 Probe design

Easy_HCR is a set of jupyter notebooks made to automate the creation of probe pairs for hybridization chain reaction (HCR). It is based on insitu_probe_generator (Kuehn *et al*., 2021). These notebooks feature automated blasting and probe pair filtering to minimize off-target effects, blasting on custom databases and probe list formatting for easy ordering from Integrated DNA Technologies, Inc (IDT). Custom database creation is necessary when Easy_HCR is used for other organisms. We recommend to design at least 20 split-initiator probe pairs per gene. Easy_HCR is available on GitHUB via https://github.com/SeuntjensLab/Easy_HCR.

Easy_HCR was used during generation of HCR v3.0 type probe pairs for fluorescent *in situ* mRNA visualization. *Ov-apolpp, Ov-ascl1, Ov-elav*, and *Ov-neuroD* were already designed as previously described in Deryckere et al., 2021 and Styfhals et al., 2022. The 33, 33, 27 and 26 split-initiator probe pairs were designed for *Ov-apolpp, Ov-ascl1, Ov-elav*, and *Ov-neuroD*, respectively (Supplementary Table T1).

DNA Oligo Pools were ordered from Integrated DNA Technologies, Inc. (probe sets are presented in Supplementary Table T1) and dissolved in Nuclease-Free Distilled Water (Invitrogen). HCR amplifiers with B1-Alexa Fluor-546, B2-Alexa Fluor-647 and B3-Alexa Fluor-488 were obtained from Molecular Instruments, Inc.

#### 2.2.2 HCR v3.0

The protocol is based upon the Molecular Instruments’ (MI) HCR v3.0 protocol for whole-mount mouse embryos (*Mus Musculus*) (Choi *et al*., 2018) with some small adaptations. Briefly, multiple octopus embryos were processed simultaneously due to their small size in 0.5 ml Eppendorf tubes. The volume of the solutions used in each step was 100 μl. During the preparation of fixed whole-mount octopus embryos, the desired amount of octopus embryos were transferred to 0.5 ml Eppendorf tubes. The embryos were thawed on ice and gradually moved to room temperature (in total half an hour). The rehydration of octopus embryos was carried out at room temperature. The octopus embryos were permeabilized by treating them for 15 minutes at room temperature using proteinase K (Roche, 10 μg/ml in PBS-DEPC). During the detection stage, the probe solutions were prepared by adding 0.4 pmol of each probe to 100 μL of probe hybridization buffer. Probes were omitted in negative controls. During the amplification stage, pre-amplification was carried out for at least 30 minutes. 3 pmol for Hairpin H1 and 3 pmol for Hairpin H2 were separately prepared (2 μL of 3 μM stock each hairpin was snap cooled: 95 °C for 90 seconds, 5 minutes on ice followed by 30 minutes at room temperature) and added to a total of 100 μl of amplification buffer. After overnight amplification, excess hairpins were removed by 3 × 100 μl 5xSSCT washes at room temperature in the dark. The embryos were incubated in 1:2000 DAPI in 5xSSCT for 2 hours followed by a 5xSSCT wash for 5 minutes. Then, embryos were transferred to the Fructose-Glycerol clearing solution described in Dekkers et al., 2019 for at least 2 days. Fructose-Glycerol clearing solution was prepared by dissolving 29,72 grams of fructose in 33 ml of glycerol and 7 ml of distilled water on a magnetic stirrer. A Refractometer was used to measure the refractive index of the Fructose-Glycerol clearing solution to validate its value being 1.45. The refractive index needs to match to the sample chamber used for imaging. A step-by-step protocol for whole mount HCR v3.0 with IHC is provided in protocol.io (dx.doi.org/10.17504/protocols.io.bxz6pp9e).

#### 2.2.3 IHC

When HCR was combined with IHC, the incubation in DAPI was skipped and the embryos were directly processed for IHC after the last excess hairpin removal wash. The whole protocol of IHC was carried out at 4°C. Embryos were incubated with the primary antibody (1:1000 rabbit anti-phospho-histone H3 (Ser10) (Millipore 06-570) for the following 2 days after the HCR protocol. Afterwards, the embryos were washed with 5xSSCT three times for 2 hours followed by adding the secondary antibody donkey anti-rabbit Alexa 488 (Life Technologies) at a final concentration of 1:300 diluted in antibody diluent (Roche) and incubated overnight. The excess secondary antibody was washed with 5xSSCT twice for 2 hours and the embryos were incubated in 1:2000 DAPI in 5xSSCT for 2 hours followed by 5xSSCT wash for 5 minutes. Fructose-Glycerol clearing was performed as described above.

### 2.3 Light sheet fluorescence microscopy imaging and analysis

Imaging was done using Zeiss Z1 Light sheet fluorescence microscopy (LSFM) (Carl Zeiss AG, Germany). The cleared and stained embryos were glued from their mantle on a metal plunger and immersed in low-viscosity immersion oil mix, as described in Deryckere et al., 2021. A refractometer was used to measure the refractive index of the immersion oil to match the refractive index of fructose-glycerol clearing solution. The z-stack coronal planes of the embryo was acquired in a series of tiles with a 20x/1.0 – refractive index 1.45 detection objective and 10x/0.2 illumination objectives. 2 μm/slice was chosen. The number of tiles was determined by considering the top-left and bottom-right coordinates and 20% tile overlap. The acquired tiles were stitched together using Arivis (Vision4D, Zeiss Edition 3.1.4). Then, image analysis such as manual reconstruction of the brain and stellate ganglia, 3D rendering and background reduction was also carried out using this software. Furthermore, fluorescent background was removed and signal-to-background ratio on light sheet images was optimized on ARIVIS software (Supplementary Figure F2).

## 3 Results and Discussion

In this study, we report for the first time on optimization of whole mount RNA multiplexed in situ hybridization chain reaction (HCR) combined with immunohistochemistry, clearing and imaging to visualize Octopus vulgaris neurogenesis. An overview of the methodology pipeline is depicted in Figure 1.

**Figure 1:**
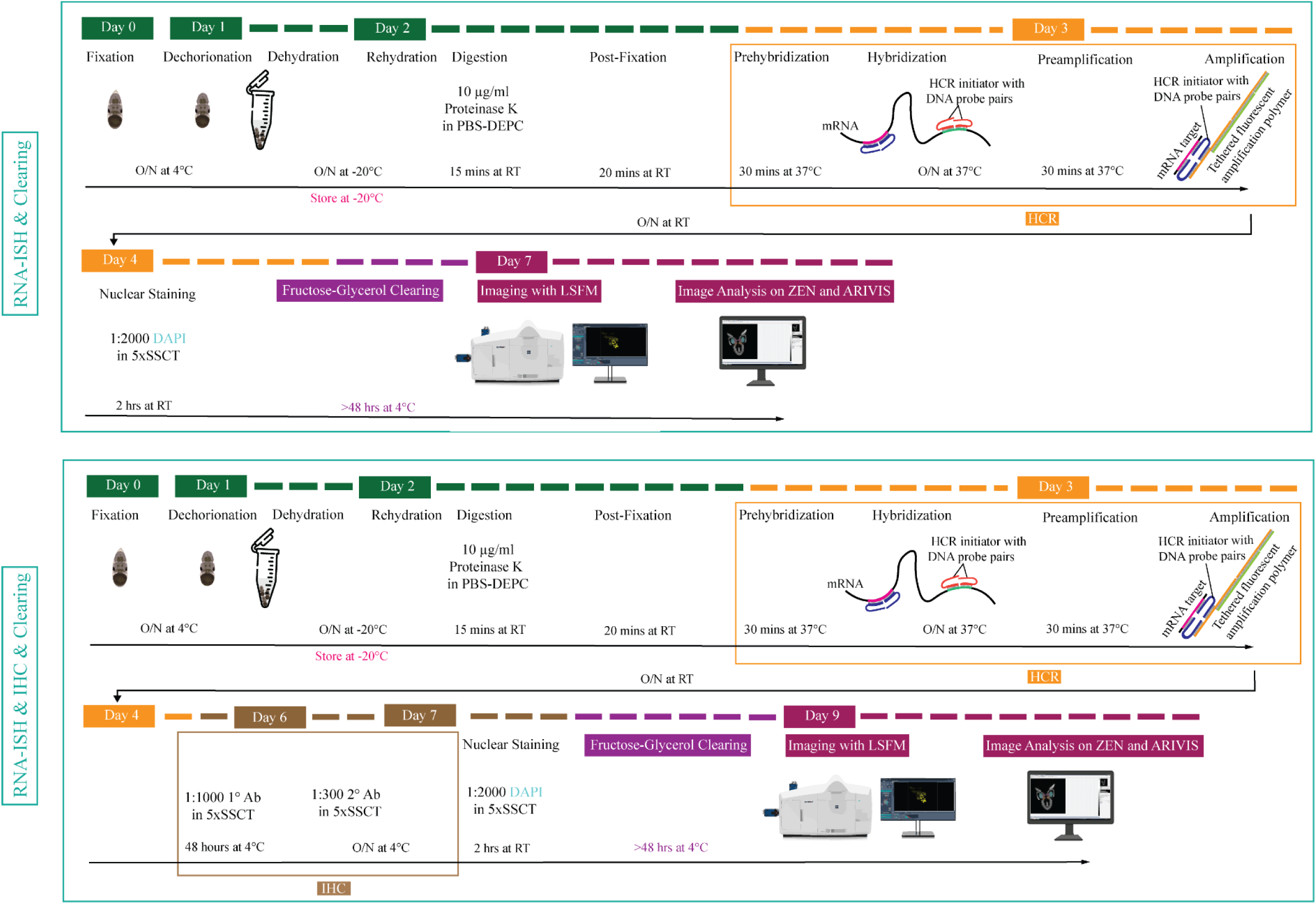
Overview of experimental pipeline for Octopus vulgaris embryos. RNA in situ hybridization chain reaction version 3.0 (RNA-ISH) and immunohistochemistry (IHC) are followed by Fructose-Glycerol clearing and imaging with Light Sheet Fluorescence Microscopy (LSFM). The final images (3D images and Z-stack planes) as well as videos are acquired, processed and analyzed with ZEN (black edition) and ARIVIS VISION4D v.3.1.4 software. For developmental stage XV embryo (its size is approximately 1,25 mm x 0,88 mm), RNA-ISH & Clearing & Imaging & Image Analysis takes approximately 7 days whereas, RNA-ISH & IHC & Clearing & Imaging & Image Analysis takes around 9 days. (This figure is designed using a resource from freepik.com).

### 3.1 Manual segmentation versus Hybridization Chain Reaction to visualize the developing nervous system

In order to benchmark our method, we first delineated the central nervous system of developmental stage XV octopus embryo using histological nuclear staining only (Figure 2). The embryo was stained by using the nuclear marker DAPI (Figure 2A-C). Afterwards, the central brain (supra-esophageal and sub-esophageal masses, and laterally located optic lobes) as well as stellate ganglia were manually segmented and reconstructed based on Marquis, 1989 (Figure 2D-I, Supplementary Video V1). Next, the developing nervous system of a developmental stage XV octopus embryo was visualized by HCR using the pan-neuronal marker *Ov-elav* (Figure 2J-L). To reduce the time of probe pair design, *in silico* validation and ordering, we developed an automated tool called Easy_HCR (available on https://github.com/SeuntjensLab/Easy_HCR) that was based on insitu_probe_generator (Kuehn *et al*., 2021). While both manual segmentation and *Ov-elav* HCR created a 3D view on the nervous system, manual segmentation was far more time-consuming and heavily dependent on the expert’s interpretation. HCR *Ov-elav* clearly delineated the neuronal cells and revealed the precise location of the gastric ganglion as well as the two buccal ganglia. Furthermore, manual segmentation is only feasible if the brain has developed to a certain point which allows experts to be able to unequivocally distinguish it from the surrounding tissues. Therefore, HCR combined with Light sheet imaging is a more accurate and less time-consuming method to visualize organ morphogenesis.

**Figure 2:**
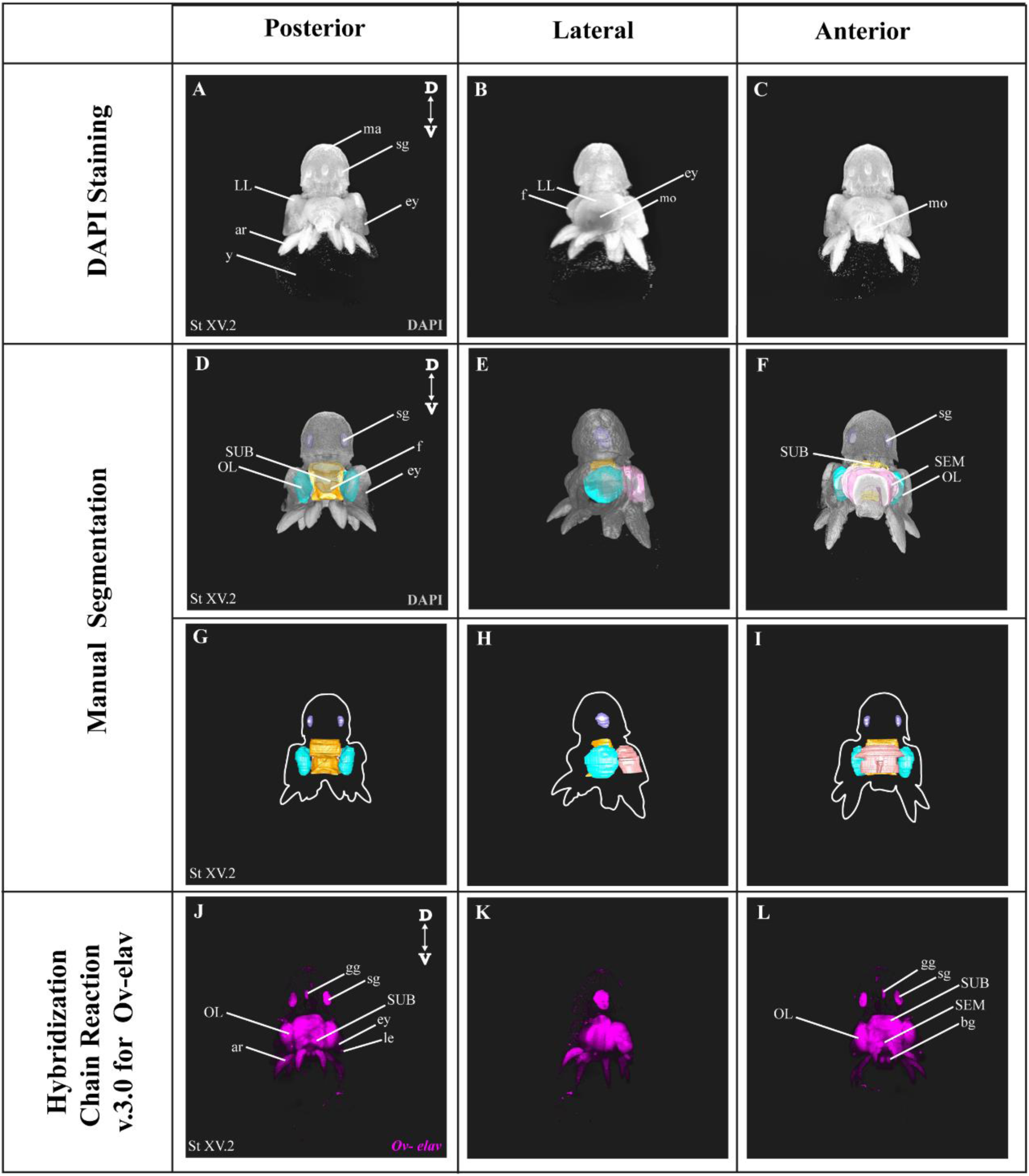
Manual Segmentation vs. *in situ* hybridization chain reaction. (A-C) Maximum intensity projection of DAPI-stained developmental stage XV octopus embryo in 3D view. (D-F) 3D volumetric octopus embryo with manually reconstructed central brain as well as stellate ganglia. *Color legend: light blue, optic lobes; pink, supraesophageal mass; orange, subesophageal mass; and dark blue, stellate ganglia*. (G-I) Manually reconstructed central brain as well as stellate ganglia. (J-L) *in situ* hybridization chain reaction (HCR) for *Ov-elav* on a developmental stage XV embryo. A stage XV octopus embryo is approximately 1,25 mm x 0,88 mm, from the top of the mantle till the end of the arms. *Abbreviations: ar, arm*; *bg, buccal ganglia*; *D, dorsal*; *ey, eye*; *fu, funnel*; *gg, gastric ganglion*; *le, lens*; *LL, lateral lip*; *ma, mantle*; *n, neuropil*; *OL, optic lobe*; *SEM, supraesophageal mass*; *sg, stellate ganglion*; *SUB, subesophageal mass*; *V, ventral; y, yolk*.

### 3.2 Multiplexing *in situ* Hybridization Chain Reaction

As a next step, we multiplexed HCR mRNA detection by combining the pan-neuronal marker (*Ov-elav*) with a glial marker (*Ov-apolpp*). As controls, we measured autofluorescence on each channel as well as performed HCR using only-hairpins without any probe conditions (Supplementary Figure F1). *Ov-elav* expression visualized the central brain masses, optic lobes, stellate, mouth and gastric ganglia as well as the neurons in the arms, while *Ov-apolpp* was mainly expressed within the neuropil located in the central brain, optic lobes and arms (Figure 3). The expression of *Ov-elav* and *Ov-apolpp* observed in whole mount octopus embryos matches the expression data seen on transverse sections (Deryckere *et al*., 2021; Styfhals *et al*., 2022). The z-stack overview and 3D view of the multiplexed HCR of *Ov-elav* and *Ov-apolpp* is provided in the Supplementary Information (Supplementary Video V2-4).

**Figure 3:**
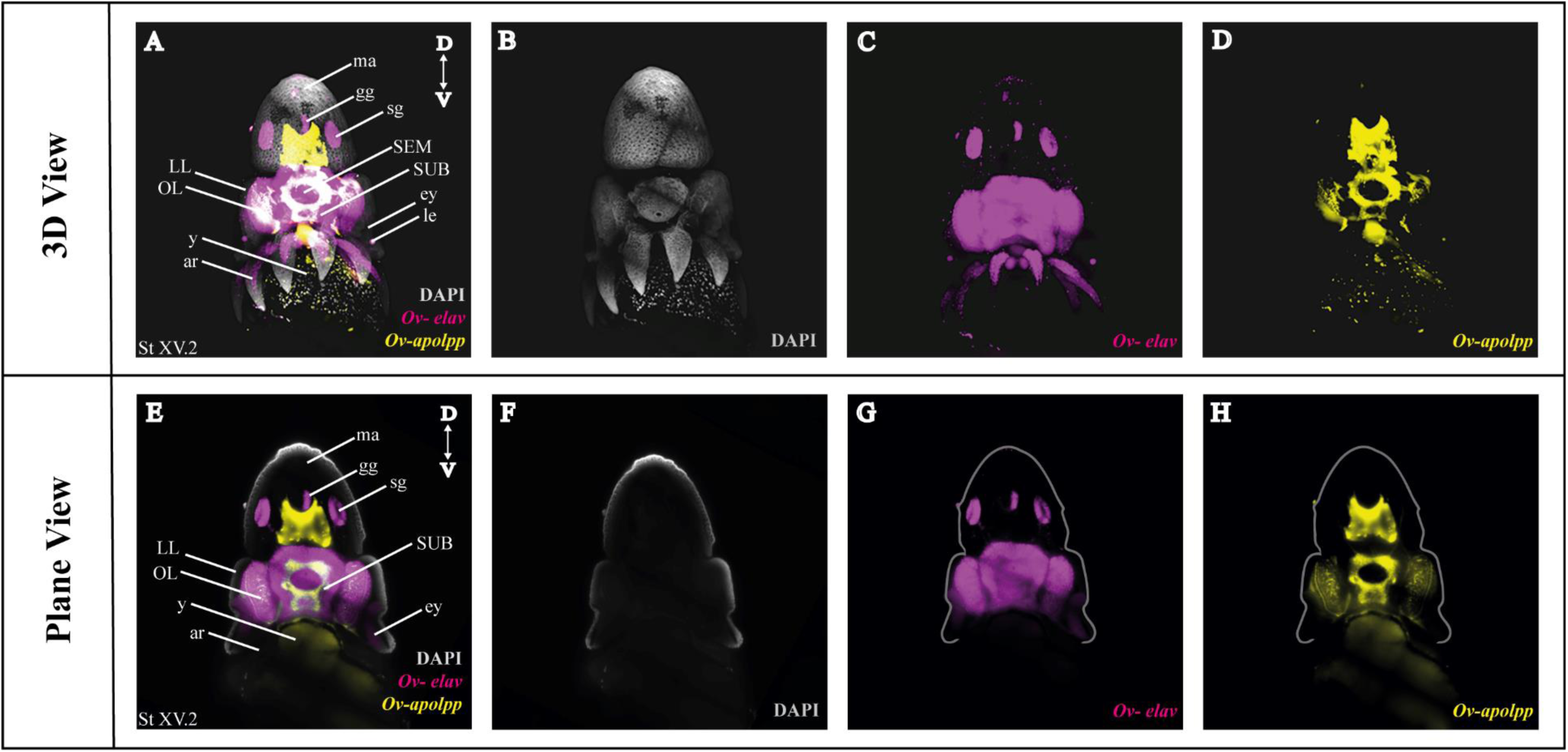
Whole Mount HCR v3.0 followed by Fructose-Glycerol Clearing on an *Octopus vulgaris* embryo (developmental stage XV) imaged with LSFM. Top panel illustrates the merged 3D view from the posterior side of the embryo, and bottom panel shows a single plane of a coronal section. (A) Overview image showing the expression of *Ov-elav and Ov-apolpp* on a Stage XV embryo in 3D. Note that only high-level expression is retained on the merged view. DAPI (in grey) is used for nuclear labelling. (B-D) 3 individual channels from A. (E) Overview image showing the expression of *Ov-elav and Ov-apolpp* on a coronal section of Stage XV embryo. (F-H) 3 individual channels from E. *Abbreviations: ar, arm*; *D, dorsal*; *ey, eye*; *fu, funnel*; *gg, gastric ganglion*; *LL, lateral lip*; *ma, mantle*; *n, neuropil*; *OL, optic lobe*; *SEM, supraesophageal mass*; *sg, stellate ganglion;SUB, subesophageal mass*; *V, ventral*; *y, yolk*.

### 3.3 Combining whole mount multiplexed *in situ* Hybridization Chain Reaction with Immunohistochemistry

After optimization of multiplexed HCR v3.0 on whole mount octopus embryos, we tested its compatibility with immunohistochemistry. Multiplexed hybridization of neuronal progenitor (*Ov-ascl1*) and neuronal precursor (*Ov-neuroD)* markers was followed by an immunostaining for the mitotic marker phosphorylated-histone H3 (PH3) (Figure 4). *Ov-ascl1* was mainly expressed in the neurogenic lateral lips and retina, while *Ov-neuroD* was mainly expressed in the transition zones connecting the neurogenic area with the central brain (Figure 4F-H). The expression of *Ov-neuroD* in the transition zone is outlined in Figure 4H. Interestingly, in 3D view, the posterior transition zone appeared to have the shape of a double bow (Supplementary Video V7). *Ov-ascl1* and *Ov-neuroD* expression observed in whole mount octopus embryo is complementary to the expression seen on transverse sections (Deryckere et al., 2021). PH3+ cells were mainly present on the skin, retina, lateral lips and arms (Figure 4E). The z-stack overview as well as 3D view of the multiplexed HCR of *Ov-ascl1* and *Ov-neuroD* with IHC of PH3 is provided in the Supplementary Information (Supplementary Video V5-7).

**Figure 4:**
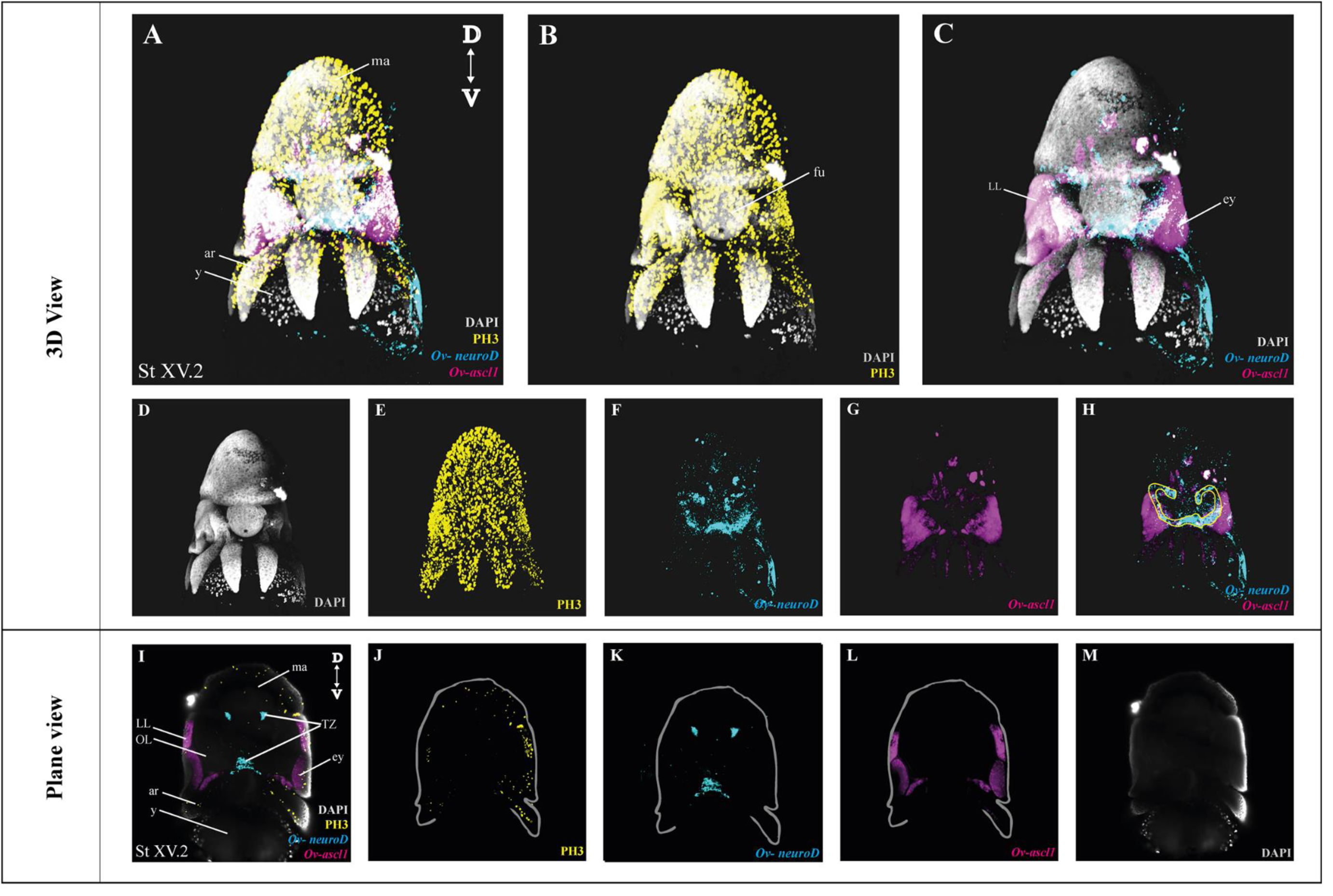
Whole Mount multiplexed HCR v3.0-IHC followed by Fructose-Glycerol Clearing on an *Octopus vulgaris* embryo (developmental stage XV) imaged with LSFM to visualize neurogenesis. (A) Overview image showing the expression of *Ov-ascl1 and Ov-neuroD* and presence of mitotic cells (PH3+) on a Stage XV embryo in 3D view. DAPI (in grey) is used for nuclear labelling. (B) Image illustrating mitotic PH3+ cells with DAPI which is an indication of successful IHC after HCR. (C) Multiplexed HCR Image of *Ov-ascl1 and Ov-neuroD* with DAPI. (D-G) Separate channels from A. (H) Overlay of *Ov-ascl1 and Ov-neuroD* show mutually exclusive expression. Yellow line indicates the transition zone area. (I) Overview image showing the expression of *Ov-ascl1 and Ov-neuroD* and presence of mitotic cells (PH3+) on a coronal section of Stage XV embryo. (J-M) 4 individual channels from I. *Abbreviations: ar, arm*; *D: dorsal*; *ey, eye*; *fu, funnel*; *LL, lateral lip*; *ma, mantle*; *OL, optic lobe*; *SUB, subesophageal mass; V: ventral; y, yolk*.

#### 3.4 Comparing different water-based clearing methods and their compatibility with HCR

Four different (CUBIC, TDE, DEEP-clear and Fructose-Glycerol) water-based clearing methods were compared for their clearing performance as well as to evaluate whether the signal intensity of HCR was retained after clearing on octopus embryos (see also Table 1) (Costantini *et al*., 2015; Nguyen, 2017; Dekkers *et al*., 2019; Pende *et al*., 2020). CUBIC (Clear, Unobstructed Brain/Body Imaging Cocktails and Computational analysis) clearing had mild clearing properties since the eye pigmentation was not completely removed which hindered brain imaging during acquisition. Furthermore, swelling of the tissue was observed. Also, no signal was observed when CUBIC was carried out after HCR on octopus embryos. TDE (60% 2,2’-thiodiethanol in PBS-DEPC) immersion was the worst-performing clearing method since no clearing of the pigmented tissue was observed even after several days of incubation. Therefore, its compatibility with HCR was not tested. The DEEP-Clear (DEpigmEntation-Plus-Clearing method) was the best to clear the octopus samples since the majority of the eye pigmentation was cleared which allowed minimal disturbance of the brain, yet it also completely wiped out the HCR signal. Fructose-Glycerol Clearing was the only option to sufficiently clear and at the same time preserve the HCR signal after clearing (Table 1).

**Table 1:** Comparison of different water-based clearing methods and their compatibility with HCR on whole mount octopus embryos and paralarvae.

| Clearing Method | Clearing Performance | Signal intensity of HCR after clearing | Observations |
| --- | --- | --- | --- |
| **CUBIC** | ++ | - | Incomplete removal of eye pigmentation which hinders the brain imaging. Tissue swelling observed. |
| **60% TDE** | + | Not tested | The pigmented tissue, especially eyes and chromatophores, is not cleared. Tissue shrinkage observed. |
| **DEEP-Clear** | +++ | - | Minimal disturbance from eye pigmentation during acquisition after clearing |
| **Fructose-Glycerol** | ++ | +++ | Pigmented tissue is cleared adequately, which allows brain imaging. Tissue shrinkage observed. |

HCR with different clearing methods has been published over the last few years. Although CUBIC clearing was not compatible with HCR in octopus embryos, positive results have been published on rat brains. It was shown that both CUBIC and CLARITY clearing could be used for clearing the rat brains to visualize *Arc* mRNA (Nguyen, 2017). PACT (passive CLARITY technique) tissue hydrogel embedding and clearing has been used after single molecule HCR in cell culture and whole-mount zebrafish embryos (Shah *et al*., 2016). HCR in combination with fructose-glycerol clearing has been used for intact tails (somites and presomitic mesoderm) of mouse embryos and afterwards, further adapted by Lütolf Group for gastruloids (3D aggregates of mouse embryonic stem cells) (Sanchez, Miyazawa and Molecular Instruments, 2019; Vianello, Park and Lutolf, 2021). Recently, glycerol (50-70%) clearing was used for a wide range of whole mount samples (Bruce *et al*., 2021). Apart from the water-based clearing methods tested, solvent-based methods, such as iDISCO+ and uDISCO, have been used successfully in combination with HCR (Kramer *et al*., 2018; Lin *et al*., 2018). The compatibility and efficiency of HCR with CLARITY and iDISCO+ in fresh-frozen rodent brain tissues and postmortem human brain blocks was previously tested (Kumar et al., 2021). An alternative for clearing methods is fluorescence tomography. For instance, combination of whole mount HCR with fluorescence tomography has been used for finding the exact location of *Cre* mRNA in a Thy1-Cre mouse brain (Guo *et al*., 2019).

## 4 Conclusion

Our aim was to report an optimized protocol for whole mount HCR v3.0 and its compatibility with IHC as well as a water-based tissue clearing method on octopus embryos for understanding neural anatomy and neurogenesis in 3D. We believe that the proposed experimental pipeline can be adapted to other model and non-model organisms. Also, HCR has a wide range of applications in various fields, such as in biomedical purposes. For instance, it can be used for studying development and creating developmental atlases for specific systems or for understanding diseases such as pathogen detection and behavior in chronic infections (Choi, Beck and Pierce, 2014; Bi, Yue and Zhang, 2017; Wu *et al*., 2021). Whole mount HCR provides a more precise 3D view compared to manual segmentation based solely on nuclear labeling. Our automated tool Easy_HCR can be used for automated probe pair design. Comparison of 4 different water-based clearing protocols should help the experimenter to pick the most robust method when performing whole mount HCR in combination with 3D imaging.

## Supporting information

Supplementary General Information

## 9 Author Contributions

AME, RS, and SM and LMa performed the experiments. AME, RS and SM analyzed and interpreted the data. ES supervised the study. AME and ES wrote the original draft of the manuscript. All authors contributed to review and editing.

## 11 Acknowledgments

The authors would like to thank Eduardo Almansa (Instituto Español de Oceanografía, Santa Cruz de Tenerife, Spain) for his support and supplying us with octopus eggs. We would like to thank Astrid Deryckere for helping us to initialize the study, and all members of the Seuntjens and Arckens labs for critical discussions.

## 12 Data Availability Statement

The original contributions presented in the study are included in the article/Supplementary Material, further inquiries can be directed to the corresponding author.

