## Supplementary Table T1 for "Optimization of Whole Mount RNA multiplexed in situ Hybridization Chain Reaction with Immunohistochemistry, Clearing and Imaging to visualize octopus neurogenesis"

**Supplementary Table T1. Nucleotide sequences of HCR probes**

| B1_OvAscl1_33_Dla2 | GAGGAGGGCAGCAAACGGaaGCGACCGAAGAAACATCTCAAGGGA |
| --- | --- |
| B1_OvAscl1_33_Dla2 | GGCCCCCTCTAATTATCACCTCTTTtaGAAGAGTCTTCCTTTACG |
| B1_OvAscl1_33_Dla2 | GAGGAGGGCAGCAAACGGaaCGCTGCTACAGAGGCCTGGTTGGAA |
| B1_OvAscl1_33_Dla2 | AATATTTCACTCCGATTCGGAAGTGtaGAAGAGTCTTCCTTTACG |
| B1_OvAscl1_33_Dla2 | GAGGAGGGCAGCAAACGGaaCTTAGACAAAAACTGAAGGACTGTT |
| B1_OvAscl1_33_Dla2 | CACCTTCCGAGTTATTTTTTTAGAGtaGAAGAGTCTTCCTTTACG |
| B1_OvAscl1_33_Dla2 | GAGGAGGGCAGCAAACGGaaAAATATATAGTTAATGATGTATGTA |
| B1_OvAscl1_33_Dla2 | TTGATATTAAATCAAAAGGTATTTAtaGAAGAGTCTTCCTTTACG |
| B1_OvAscl1_33_Dla2 | GAGGAGGGCAGCAAACGGaaTGTTTATTTAATATAAATTGGCGCA |
| B1_OvAscl1_33_Dla2 | ATTTATAAATACATATGTTCGACTGtaGAAGAGTCTTCCTTTACG |
| B1_OvAscl1_33_Dla2 | GAGGAGGGCAGCAAACGGaaCAGTGTCTGTGCGTTTGCGTATTCT |
| B1_OvAscl1_33_Dla2 | CCATATACACTTGTTACGTATGAAGtaGAAGAGTCTTCCTTTACG |
| B1_OvAscl1_33_Dla2 | GAGGAGGGCAGCAAACGGaaATAGTTTGTACAAAAATGTTGAGAT |
| B1_OvAscl1_33_Dla2 | AAAATATATTATATTATATGGAAGAtaGAAGAGTCTTCCTTTACG |
| B1_OvAscl1_33_Dla2 | GAGGAGGGCAGCAAACGGaaTAAATTTGTCACTCTCTGTTCTACA |
| B1_OvAscl1_33_Dla2 | GCACTGAACTTCATGTATGGTTGTAtaGAAGAGTCTTCCTTTACG |
| B1_OvAscl1_33_Dla2 | GAGGAGGGCAGCAAACGGaaTATTAATTCTCCACCCGTTCCCACC |
| B1_OvAscl1_33_Dla2 | AAACTAAACTGGGATGACATTTTTAtaGAAGAGTCTTCCTTTACG |
| B1_OvAscl1_33_Dla2 | GAGGAGGGCAGCAAACGGaaCTAGTTGAGGGTACAGTTTATTCTA |
| B1_OvAscl1_33_Dla2 | CAAAAATAAAAGTCTCAAGATTTCAtaGAAGAGTCTTCCTTTACG |
| B1_OvAscl1_33_Dla2 | GAGGAGGGCAGCAAACGGaaAAATCTATTCACAAGTAGTAGATGA |
| B1_OvAscl1_33_Dla2 | ATCTTAGCCTTGTGAATTGGCCATAtaGAAGAGTCTTCCTTTACG |
| B1_OvAscl1_33_Dla2 | GAGGAGGGCAGCAAACGGaaGATGAGCAATGGATGATTTAACAAT |
| B1_OvAscl1_33_Dla2 | ATGTCAGTGGATTAGCAAACTGTATtaGAAGAGTCTTCCTTTACG |
| B1_OvAscl1_33_Dla2 | GAGGAGGGCAGCAAACGGaaCAAAATTTCGCGGGCAATTATTCAT |
| B1_OvAscl1_33_Dla2 | TACTATTGTCAGTTTATGTTCTGCAtaGAAGAGTCTTCCTTTACG |
| B1_OvAscl1_33_Dla2 | GAGGAGGGCAGCAAACGGaaATATAAATTATGCCATTTTTGAAAT |
| B1_OvAscl1_33_Dla2 | TACCTCCAATCCTAACAGATGAAGTtaGAAGAGTCTTCCTTTACG |
| B1_OvAscl1_33_Dla2 | GAGGAGGGCAGCAAACGGaaTAGAATTCCTTATGATTTTTATTTG |
| B1_OvAscl1_33_Dla2 | TGTATGACCATGTTTTCGAGCATATtaGAAGAGTCTTCCTTTACG |
| B1_OvAscl1_33_Dla2 | GAGGAGGGCAGCAAACGGaaAATTCCGTGAAACCGATCGGCAAAA |
| B1_OvAscl1_33_Dla2 | TATATTTTCTTATGCGTTAAAAACAtaGAAGAGTCTTCCTTTACG |
| B1_OvAscl1_33_Dla2 | GAGGAGGGCAGCAAACGGaaAAGTTCGTGTGCTGTTGGAAGAAGA |
| B1_OvAscl1_33_Dla2 | GTTTCAGCGACAGTCATGTGTTTGGtaGAAGAGTCTTCCTTTACG |
| B1_OvAscl1_33_Dla2 | GAGGAGGGCAGCAAACGGaaTCAGAAGGCGATACGATATACGTAT |
| B1_OvAscl1_33_Dla2 | ATTTCACGTTTCGGAACTGACCCTTtaGAAGAGTCTTCCTTTACG |
| B1_OvAscl1_33_Dla2 | GAGGAGGGCAGCAAACGGaaTCATTTTAAAGAAAATAAACTACTA |
| B1_OvAscl1_33_Dla2 | ATGTTCAACAGATATATTTTATATAtaGAAGAGTCTTCCTTTACG |
| B1_OvAscl1_33_Dla2 | GAGGAGGGCAGCAAACGGaaTAATTTAGTTTTTATATTTCTTTTC |
| B1_OvAscl1_33_Dla2 | TCTTTACCAAGAAAATAAACTTCTAtaGAAGAGTCTTCCTTTACG |
| B1_OvAscl1_33_Dla2 | GAGGAGGGCAGCAAACGGaaCTTTTGCTTTGGAATTCAGATTCTA |
| B1_OvAscl1_33_Dla2 | GTTTTTAAACATTAGTAAATATATTtaGAAGAGTCTTCCTTTACG |
| B1_OvAscl1_33_Dla2 | GAGGAGGGCAGCAAACGGaaACGGAATGACAGCGAGTTTTCAATT |
| B1_OvAscl1_33_Dla2 | TTGGTTCTGTAGAGAAACTTTCCATtaGAAGAGTCTTCCTTTACG |
| B1_OvAscl1_33_Dla2 | GAGGAGGGCAGCAAACGGaaCCTCTGCAGCGATTTCCTGACATTT |
| B1_OvAscl1_33_Dla2 | TTTTTCGTTATGAACATGTGGCTGTtaGAAGAGTCTTCCTTTACG |
| B1_OvAscl1_33_Dla2 | GAGGAGGGCAGCAAACGGaaTTATTTCAGGTCTTTGTTGTTTTGT |
| B1_OvAscl1_33_Dla2 | TTCCTTTTTATTTTCATATGATTTTtaGAAGAGTCTTCCTTTACG |
| B1_OvAscl1_33_Dla2 | GAGGAGGGCAGCAAACGGaaTATAGAACTTGTTTCTTTTACTTTT |
| B1_OvAscl1_33_Dla2 | TATATACGCATGCATGAATGCACATtaGAAGAGTCTTCCTTTACG |
| B1_OvAscl1_33_Dla2 | GAGGAGGGCAGCAAACGGaaGAGCAAGAACTCGGTCGCGGCGAGT |
| B1_OvAscl1_33_Dla2 | GCACTAAGTCCTTCATAAGAACTATtaGAAGAGTCTTCCTTTACG |
| B1_OvAscl1_33_Dla2 | GAGGAGGGCAGCAAACGGaaGTTGACGGTGTCACTATCCTCAACA |
| B1_OvAscl1_33_Dla2 | GGTCAAACAGCTACTATCGAAGATAtaGAAGAGTCTTCCTTTACG |
| B1_OvAscl1_33_Dla2 | GAGGAGGGCAGCAAACGGaaCGTTCGGTATATGTTCTCTAAGCGT |
| B1_OvAscl1_33_Dla2 | TTTTTTTGTTCTTACTGCCCTTCGGtaGAAGAGTCTTCCTTTACG |
| B1_OvAscl1_33_Dla2 | GAGGAGGGCAGCAAACGGaaGGCAAGTAGTTGGTGTGGTTGAAAT |
| B1_OvAscl1_33_Dla2 | ACGGCAACTGTCTGTGGTCGATGCAtaGAAGAGTCTTCCTTTACG |
| B1_OvAscl1_33_Dla2 | GAGGAGGGCAGCAAACGGaaGTGCGTTCTCTTTCATCGATATCGT |
| B1_OvAscl1_33_Dla2 | GACGTCGCTTACAGTTCATCAGCTCtaGAAGAGTCTTCCTTTACG |
| B1_OvAscl1_33_Dla2 | GAGGAGGGCAGCAAACGGaaCGTCGTTGTGGAGTTTGTGGTCGCC |
| B1_OvAscl1_33_Dla2 | GACTGATTTCGAATTGGGAGTCGTCtaGAAGAGTCTTCCTTTACG |
| B1_OvAscl1_33_Dla2 | GAGGAGGGCAGCAAACGGaaGTTGCCGCCACTGCCGTGGTCGTTG |
| B1_OvAscl1_33_Dla2 | GTCGTCGTCGTCGTTGGAAACCCGTtaGAAGAGTCTTCCTTTACG |
| B1_OvAscl1_33_Dla2 | GAGGAGGGCAGCAAACGGaaATTGTATTTCGTGCCGCTGCTTGTT |
| B1_OvAscl1_33_Dla2 | ATTGGTTCCGGCACCAGTGCCGCTGtaGAAGAGTCTTCCTTTACG |
| Ov-Apolpp_B3_33PP | GTCCCTGCCTCTATATCTTTAGGTGGATCAATAAAAAGATATATA |
| Ov-Apolpp_B3_33PP | ATTTGAGAGAGGAACAGACAGGTAGTTCCACTCAACTTTAACCCG |
| Ov-Apolpp_B3_33PP | GTCCCTGCCTCTATATCTTTTTTCCATGGTGCTCTGGTCCCCTTC |
| Ov-Apolpp_B3_33PP | TCCTCGAAGGGGAGATGATCATGTATTCCACTCAACTTTAACCCG |
| Ov-Apolpp_B3_33PP | GTCCCTGCCTCTATATCTTTATATTGAATTCCCGAGCCATAGGGT |
| Ov-Apolpp_B3_33PP | ACATTCTTCAGCGGACTCCGCATCGTTCCACTCAACTTTAACCCG |
| Ov-Apolpp_B3_33PP | GTCCCTGCCTCTATATCTTTGGCCCGTAGCTCCAGGTTCACACCA |
| Ov-Apolpp_B3_33PP | TTCAATCCCCAAGTTGGCTGGACTTTTCCACTCAACTTTAACCCG |
| Ov-Apolpp_B3_33PP | GTCCCTGCCTCTATATCTTTTTTCTGCGGATTTCTTCGGATCGTG |
| Ov-Apolpp_B3_33PP | TGTTCTCCCTCATAAGTCCAAACGCTTCCACTCAACTTTAACCCG |
| Ov-Apolpp_B3_33PP | GTCCCTGCCTCTATATCTTTGCTGGCATAAGATTGATCGCAACAT |
| Ov-Apolpp_B3_33PP | TCGTTGGTCCACTGGATTATCTTTTTTCCACTCAACTTTAACCCG |
| Ov-Apolpp_B3_33PP | GTCCCTGCCTCTATATCTTTGGCCACCATAAACGGTCGCGTTGTC |
| Ov-Apolpp_B3_33PP | GTCGTCTCGATCCGCATCCCATAGGTTCCACTCAACTTTAACCCG |
| Ov-Apolpp_B3_33PP | GTCCCTGCCTCTATATCTTTTGGCAAAAGTGATCTTTTTGTCCTT |
| Ov-Apolpp_B3_33PP | GGTAAACCTCCCGTCGGATGCCACCTTCCACTCAACTTTAACCCG |
| Ov-Apolpp_B3_33PP | GTCCCTGCCTCTATATCTTTGGGTGGTTGATTGTCACTTCAAGCT |
| Ov-Apolpp_B3_33PP | TCAACAGAGAAACCAAAACTGCGAGTTCCACTCAACTTTAACCCG |
| Ov-Apolpp_B3_33PP | GTCCCTGCCTCTATATCTTTGGCAAGATACTTATCTTTTAGGGAT |
| Ov-Apolpp_B3_33PP | ATCTTTGTCCCAGTTGCATTTGATTTTCCACTCAACTTTAACCCG |
| Ov-Apolpp_B3_33PP | GTCCCTGCCTCTATATCTTTCTTTCCCAGGCATCACTTCCTCAGG |
| Ov-Apolpp_B3_33PP | TGACTTGTTTGAAGTTGGCGTCGAATTCCACTCAACTTTAACCCG |
| Ov-Apolpp_B3_33PP | GTCCCTGCCTCTATATCTTTCGGGTCACGACCACGTCTGCCTTGT |
| Ov-Apolpp_B3_33PP | AAGATGTACTTCTTCGATTTGTTCTTTCCACTCAACTTTAACCCG |
| Ov-Apolpp_B3_33PP | GTCCCTGCCTCTATATCTTTGTTTTGACGATGAGTTTCAAATCGG |
| Ov-Apolpp_B3_33PP | TTCACTGTTCTTTTATTGCTCCATGTTCCACTCAACTTTAACCCG |
| Ov-Apolpp_B3_33PP | GTCCCTGCCTCTATATCTTTGGCCATTATCTCAAAATTGATCTCC |
| Ov-Apolpp_B3_33PP | TAACTTTATGATGAAAACTTCACGCTTCCACTCAACTTTAACCCG |
| Ov-Apolpp_B3_33PP | GTCCCTGCCTCTATATCTTTCAGACAAAATATCAAAAGAACTGAC |
| Ov-Apolpp_B3_33PP | TGAGTTTCACTTCTCCTGATCCGGTTTCCACTCAACTTTAACCCG |
| Ov-Apolpp_B3_33PP | GTCCCTGCCTCTATATCTTTTCCCCCTTTTTATTTTTGCCCATTC |
| Ov-Apolpp_B3_33PP | TTACTATATGAGGCCCTGAATTTGATTCCACTCAACTTTAACCCG |
| Ov-Apolpp_B3_33PP | GTCCCTGCCTCTATATCTTTAGTAATTACATAAGGAACATCGTTG |
| Ov-Apolpp_B3_33PP | GTCAAATCGTAATTCCGCCTTCTGGTTCCACTCAACTTTAACCCG |
| Ov-Apolpp_B3_33PP | GTCCCTGCCTCTATATCTTTAAACCGTAACAAGAAGATTCGTGAA |
| Ov-Apolpp_B3_33PP | AGAAGCTGTGACTTGGAGTTGGAGTTTCCACTCAACTTTAACCCG |
| Ov-Apolpp_B3_33PP | GTCCCTGCCTCTATATCTTTTTAGAAAGTACTCTAGATGAGATCA |
| Ov-Apolpp_B3_33PP | ATCTGCAAATGGTTTTTCTCTTTTGTTCCACTCAACTTTAACCCG |
| Ov-Apolpp_B3_33PP | GTCCCTGCCTCTATATCTTTATCAAATTCATAGTTGCCAGATATG |
| Ov-Apolpp_B3_33PP | CACATGGATCATGAGGTCCCCAACATTCCACTCAACTTTAACCCG |
| Ov-Apolpp_B3_33PP | GTCCCTGCCTCTATATCTTTCAACGTCTTTCTTCTTTCGATAGAC |
| Ov-Apolpp_B3_33PP | ATATATTCATTTTGCTGGTTATCTTTTCCACTCAACTTTAACCCG |
| Ov-Apolpp_B3_33PP | GTCCCTGCCTCTATATCTTTAAAGAACTTTCAATTGTTATATTTG |
| Ov-Apolpp_B3_33PP | TCTAATGATTTTGATGCTTCGAATTTTCCACTCAACTTTAACCCG |
| Ov-Apolpp_B3_33PP | GTCCCTGCCTCTATATCTTTGACTTCCCCAGTGAACAAAGCCATT |
| Ov-Apolpp_B3_33PP | CAGCTTGACATCTGTAGTAATGGTGTTCCACTCAACTTTAACCCG |
| Ov-Apolpp_B3_33PP | GTCCCTGCCTCTATATCTTTATTTTCCATCTATGTTGTCTTTGAC |
| Ov-Apolpp_B3_33PP | TCAGTTTATCATTGTAATGGACACTTTCCACTCAACTTTAACCCG |
| Ov-Apolpp_B3_33PP | GTCCCTGCCTCTATATCTTTAAAATATCTTTGTTGCCTACTGTTA |
| Ov-Apolpp_B3_33PP | TTTTTCACGACAGAATACATTGAGTTTCCACTCAACTTTAACCCG |
| Ov-Apolpp_B3_33PP | GTCCCTGCCTCTATATCTTTTCTTGTTATTAACCAGTTTATAATG |
| Ov-Apolpp_B3_33PP | TCATTTCAGCTAAAAACTTAAACATTTCCACTCAACTTTAACCCG |
| Ov-Apolpp_B3_33PP | GTCCCTGCCTCTATATCTTTCTATAGAAGTCTTTGAATTTCATTT |
| Ov-Apolpp_B3_33PP | TTATAATACCATTTCCAAGTTGCCATTCCACTCAACTTTAACCCG |
| Ov-Apolpp_B3_33PP | GTCCCTGCCTCTATATCTTTTTCGAGTGTTGCCACAGCAGAAGTG |
| Ov-Apolpp_B3_33PP | TTCGAAACCCTCCCAAGGCAGGAACTTCCACTCAACTTTAACCCG |
| Ov-Apolpp_B3_33PP | GTCCCTGCCTCTATATCTTTCGGATTCGTGTTTCAAAGAAATATC |
| Ov-Apolpp_B3_33PP | CCCAGCCAACCAAAAAGCTAGCATCTTCCACTCAACTTTAACCCG |
| Ov-Apolpp_B3_33PP | GTCCCTGCCTCTATATCTTTTGTTCTTTATGGAAATCACTCCACC |
| Ov-Apolpp_B3_33PP | TCTGCTTCCGATTCGGCAATAACAGTTCCACTCAACTTTAACCCG |
| Ov-Apolpp_B3_33PP | GTCCCTGCCTCTATATCTTTTGTTCTTTCTTCAATTGGTTACTGA |
| Ov-Apolpp_B3_33PP | GACAACGTGTTCTCGTAGTTACGAATTCCACTCAACTTTAACCCG |
| Ov-Apolpp_B3_33PP | GTCCCTGCCTCTATATCTTTTCCTTCTCTCTGAACATTGTGCTTG |
| Ov-Apolpp_B3_33PP | TCCGATTCCGCCGTACTTTTTGGCGTTCCACTCAACTTTAACCCG |
| Ov-Apolpp_B3_33PP | GTCCCTGCCTCTATATCTTTTGTGATCTAGATCCAACTCAAAATC |
| Ov-Apolpp_B3_33PP | TGTTCAGAAGAAAATTGTCTTGGTGTTCCACTCAACTTTAACCCG |
| B1_OvElav_27_Dla2 | GAGGAGGGCAGCAAACGGaaGTTCACTCAATTTGAAAGTAGAAAT |
| B1_OvElav_27_Dla2 | TAGTAAACTTATATGGACTGTCACAtaGAAGAGTCTTCCTTTACG |
| B1_OvElav_27_Dla2 | GAGGAGGGCAGCAAACGGaaGATTAAACTAAAAGACAGAATTACT |
| B1_OvElav_27_Dla2 | TACACAAATAACAGATGTAAAAGTAtaGAAGAGTCTTCCTTTACG |
| B1_OvElav_27_Dla2 | GAGGAGGGCAGCAAACGGaaAATGGCAAACAAACTATTAGTGAAA |
| B1_OvElav_27_Dla2 | AAAGCAGCTTTACTAAGAAATAAATtaGAAGAGTCTTCCTTTACG |
| B1_OvElav_27_Dla2 | GAGGAGGGCAGCAAACGGaaAAACCAGACAAAATAATGAAAAAAA |
| B1_OvElav_27_Dla2 | CAGCATGCAATGAAAAAAAAAAGCAtaGAAGAGTCTTCCTTTACG |
| B1_OvElav_27_Dla2 | GAGGAGGGCAGCAAACGGaaACAACTAATGATGCAATACTTTGGA |
| B1_OvElav_27_Dla2 | TATTAACAGAATGAAACTGTAATACtaGAAGAGTCTTCCTTTACG |
| B1_OvElav_27_Dla2 | GAGGAGGGCAGCAAACGGaaAAACAAAAAATACCAATTTCTGCCC |
| B1_OvElav_27_Dla2 | CAATAAATAATGCAAAAGAAAATAAtaGAAGAGTCTTCCTTTACG |
| B1_OvElav_27_Dla2 | GAGGAGGGCAGCAAACGGaaAAGTGTCATTAAATAAGTCAAAATA |
| B1_OvElav_27_Dla2 | TGCTTGACTAAGTTAAACTATGGGAtaGAAGAGTCTTCCTTTACG |
| B1_OvElav_27_Dla2 | GAGGAGGGCAGCAAACGGaaTGATATGCCTTATATGGCGGATGAC |
| B1_OvElav_27_Dla2 | ATATACTTTGGTGTTTTGATTTAAAtaGAAGAGTCTTCCTTTACG |
| B1_OvElav_27_Dla2 | GAGGAGGGCAGCAAACGGaaAAAAAAGTCCGCATCTATGACCACA |
| B1_OvElav_27_Dla2 | AGGCCATATGGCATGGAATACACCAtaGAAGAGTCTTCCTTTACG |
| B1_OvElav_27_Dla2 | GAGGAGGGCAGCAAACGGaaTTCTCCTTCATGTCTTAGGATTTAC |
| B1_OvElav_27_Dla2 | CTGACTAGGAGTCCATCAGGATATTtaGAAGAGTCTTCCTTTACG |
| B1_OvElav_27_Dla2 | GAGGAGGGCAGCAAACGGaaCCCGTGTGCCGAGTAAGAATCCATT |
| B1_OvElav_27_Dla2 | TATTCGTTTTGAATGATACCTGTAGtaGAAGAGTCTTCCTTTACG |
| B1_OvElav_27_Dla2 | GAGGAGGGCAGCAAACGGaaATAATTGGTCATTGTAACAAACCCA |
| B1_OvElav_27_Dla2 | TGTCTGTATAGCCATCAATGCCTCTtaGAAGAGTCTTCCTTTACG |
| B1_OvElav_27_Dla2 | GAGGAGGGCAGCAAACGGaaCGAATCACTTTCACATTTTGCACAG |
| B1_OvElav_27_Dla2 | CCTTTGCATTTCTGTGTTTGAAAATtaGAAGAGTCTTCCTTTACG |
| B1_OvElav_27_Dla2 | GAGGAGGGCAGCAAACGGaaCATCATCCTCAGTGTCTGGTGCAAG |
| B1_OvElav_27_Dla2 | CAAATGGTCCAAAGAGACGCCAAAGtaGAAGAGTCTTCCTTTACG |
| B1_OvElav_27_Dla2 | GAGGAGGGCAGCAAACGGaaATTTAGAGCATTCCCTGGAAGTATT |
| B1_OvElav_27_Dla2 | GTATACAAAAATACACCAACCAGTGtaGAAGAGTCTTCCTTTACG |
| B1_OvElav_27_Dla2 | GAGGAGGGCAGCAAACGGaaGAGTATCTGAATCTTCCAGCAGCAT |
| B1_OvElav_27_Dla2 | CCTGGCAGTAGACTAGCTTCCAATGtaGAAGAGTCTTCCTTTACG |
| B1_OvElav_27_Dla2 | GAGGAGGGCAGCAAACGGaaAGAGTTAGCAAACTTTACAGTAATA |
| B1_OvElav_27_Dla2 | AGGTACAGCATTTTTATTTGAGCTTtaGAAGAGTCTTCCTTTACG |
| B1_OvElav_27_Dla2 | GAGGAGGGCAGCAAACGGaaTGTAATTTTTGGATAGCTCTTTCAG |
| B1_OvElav_27_Dla2 | TCAGTTGCTCCTTCAGGAATGGTACtaGAAGAGTCTTCCTTTACG |
| B1_OvElav_27_Dla2 | GAGGAGGGCAGCAAACGGaaCAACACCTTTGGATAAGCCTGTGTT |
| B1_OvElav_27_Dla2 | CGATTCGTTGATCAAATCGGATGAAtaGAAGAGTCTTCCTTTACG |
| B1_OvElav_27_Dla2 | GAGGAGGGCAGCAAACGGaaGAATCCACACTGAGAAAACAATTTC |
| B1_OvElav_27_Dla2 | GTCATAAAGAATTCTGGATGTTATAtaGAAGAGTCTTCCTTTACG |
| B1_OvElav_27_Dla2 | GAGGAGGGCAGCAAACGGaaAAGCCACTTATGTACAAGTTTGCCC |
| B1_OvElav_27_Dla2 | AAGTCCAACTGAGTGAAAGACTTTGtaGAAGAGTCTTCCTTTACG |
| B1_OvElav_27_Dla2 | GAGGAGGGCAGCAAACGGaaCTTTTCTGCATCACTAGGGTATTTG |
| B1_OvElav_27_Dla2 | TCTCAATCCATTTAAAGTATTGATTtaGAAGAGTCTTCCTTTACG |
| B1_OvElav_27_Dla2 | GAGGAGGGCAGCAAACGGaaCATTTTTCCGCATCGGTTGATGCAT |
| B1_OvElav_27_Dla2 | TTAACAAAGCCATAACCTAAACTCTtaGAAGAGTCTTCCTTTACG |
| B1_OvElav_27_Dla2 | GAGGAGGGCAGCAAACGGaaAGTCAGTGTTAGCTGTTTCCGGCAG |
| B1_OvElav_27_Dla2 | GTGCTGCCGGAAGCAGGTCACAGTGtaGAAGAGTCTTCCTTTACG |
| B2_OvNeuroD_26_Dla1 | CCTCGTAAATCCTCATCAaaCTCTTTAGATAATGTCTCGAATATT |
| B2_OvNeuroD_26_Dla1 | ACAGCATAAAACCAAAAGTCGTTTCaaATCATCCAGTAAACCGCC |
| B2_OvNeuroD_26_Dla1 | CCTCGTAAATCCTCATCAaaTCCAATAAGAAGCAACGTTATAAAA |
| B2_OvNeuroD_26_Dla1 | ATCATATTTGTACGTATATTTATTCaaATCATCCAGTAAACCGCC |
| B2_OvNeuroD_26_Dla1 | CCTCGTAAATCCTCATCAaaTTAATCAAAACCAAATGTGTTATAA |
| B2_OvNeuroD_26_Dla1 | TCCGCTTCGATCAGAAATCTTTTACaaATCATCCAGTAAACCGCC |
| B2_OvNeuroD_26_Dla1 | CCTCGTAAATCCTCATCAaaAAGACAGATCATTGCAACAAAACAT |
| B2_OvNeuroD_26_Dla1 | GCCCAAACTCTGGTTACTAACCACAaaATCATCCAGTAAACCGCC |
| B2_OvNeuroD_26_Dla1 | CCTCGTAAATCCTCATCAaaAGTGATTTCAAATATACCTCGGTTA |
| B2_OvNeuroD_26_Dla1 | TGATGATAGAATCTCAGCAGCTCAAaaATCATCCAGTAAACCGCC |
| B2_OvNeuroD_26_Dla1 | CCTCGTAAATCCTCATCAaaACTGGTGGGTCACTCTGAAAATCGG |
| B2_OvNeuroD_26_Dla1 | TTTATGATATTTAAATTGTGCTCCAaaATCATCCAGTAAACCGCC |
| B2_OvNeuroD_26_Dla1 | CCTCGTAAATCCTCATCAaaGCTGAGGTTGAGCACAACCATTGTA |
| B2_OvNeuroD_26_Dla1 | TATCTTCTAATAAGACATAGGGGCTaaATCATCCAGTAAACCGCC |
| B2_OvNeuroD_26_Dla1 | CCTCGTAAATCCTCATCAaaGTACGTGCTGTTTGAGTTGTTGTTA |
| B2_OvNeuroD_26_Dla1 | ACGCCTGATACCAACATCATTGACCaaATCATCCAGTAAACCGCC |
| B2_OvNeuroD_26_Dla1 | CCTCGTAAATCCTCATCAaaGTGTTGTTTGATGTCAAACCGGGTG |
| B2_OvNeuroD_26_Dla1 | TTATTGTTGGTGTTATAACGAAGATaaATCATCCAGTAAACCGCC |
| B2_OvNeuroD_26_Dla1 | CCTCGTAAATCCTCATCAaaTATGTTCTGCATTCAACGGACTCAG |
| B2_OvNeuroD_26_Dla1 | TACTTCCAGTAGGCAACGAGGCTTCaaATCATCCAGTAAACCGCC |
| B2_OvNeuroD_26_Dla1 | CCTCGTAAATCCTCATCAaaTTGATGGGTGGGCGGTGGTAGTGAA |
| B2_OvNeuroD_26_Dla1 | ACGCGGTGTAGTGTAATTAGTGTACaaATCATCCAGTAAACCGCC |
| B2_OvNeuroD_26_Dla1 | CCTCGTAAATCCTCATCAaaGAAGAAGACAATCGATTGGATGGAA |
| B2_OvNeuroD_26_Dla1 | ATCATGCCGTTAGATGGGGCGTGAGaaATCATCCAGTAAACCGCC |
| B2_OvNeuroD_26_Dla1 | CCTCGTAAATCCTCATCAaaTCATGTTTGGTTGTGGCTGTTGTGA |
| B2_OvNeuroD_26_Dla1 | GGCCTACCATTGTGATTGGCCCGATaaATCATCCAGTAAACCGCC |
| B2_OvNeuroD_26_Dla1 | CCTCGTAAATCCTCATCAaaGCTGCTGTTCCTGCGGTTGCATACT |
| B2_OvNeuroD_26_Dla1 | GGGGGTGAGGAAGTGGTGGAATGTGaaATCATCCAGTAAACCGCC |
| B2_OvNeuroD_26_Dla1 | CCTCGTAAATCCTCATCAaaGGAGGATATGGAATTTGAGTAACAG |
| B2_OvNeuroD_26_Dla1 | TGATGCAGAGTCGTTGCATAGGAAGaaATCATCCAGTAAACCGCC |
| B2_OvNeuroD_26_Dla1 | CCTCGTAAATCCTCATCAaaCTGAAGACATCCAGCAACGAGATTC |
| B2_OvNeuroD_26_Dla1 | ATCGGGGTGCAATGTTCGAGGATTAaaATCATCCAGTAAACCGCC |
| B2_OvNeuroD_26_Dla1 | CCTCGTAAATCCTCATCAaaGCTTTCGCAAAACTGACACCATCTG |
| B2_OvNeuroD_26_Dla1 | GTGTTTTGAGAAAGACCTTTTGACAaaATCATCCAGTAAACCGCC |
| B2_OvNeuroD_26_Dla1 | CCTCGTAAATCCTCATCAaaCCATGCATGCGGTTGCGTTCTCGTG |
| B2_OvNeuroD_26_Dla1 | CGCAAAGTGTCCAATGCATCGTTCAaaATCATCCAGTAAACCGCC |
| B2_OvNeuroD_26_Dla1 | CCTCGTAAATCCTCATCAaaACGTGGTTCGTTCTTGTCTTTGGAA |
| B2_OvNeuroD_26_Dla1 | CTTCTTTGGACCACGTTTCTTTGGAaaATCATCCAGTAAACCGCC |
| B2_OvNeuroD_26_Dla1 | CCTCGTAAATCCTCATCAaaGTCATCTTCTGAACATGTGTCATCT |
| B2_OvNeuroD_26_Dla1 | TTCTGGTTCGCTGCCATAATCATCCaaATCATCCAGTAAACCGCC |
| B2_OvNeuroD_26_Dla1 | CCTCGTAAATCCTCATCAaaTGAACATCTGCATTCCGCTCGTAGT |
| B2_OvNeuroD_26_Dla1 | CTGTTTCCAAACTTTTGATAGGCATaaATCATCCAGTAAACCGCC |
| B2_OvNeuroD_26_Dla1 | CCTCGTAAATCCTCATCAaaTCGATGTATCTAAGCAATCATGCTT |
| B2_OvNeuroD_26_Dla1 | TTCAGTTTTAAACCCGTTTAAACGGaaATCATCCAGTAAACCGCC |
| B2_OvNeuroD_26_Dla1 | CCTCGTAAATCCTCATCAaaGACATAAATAAATCTGGCTTGACAA |
| B2_OvNeuroD_26_Dla1 | GATGAAGTTCCAGTCTGAATGAAGTaaATCATCCAGTAAACCGCC |
| B2_OvNeuroD_26_Dla1 | CCTCGTAAATCCTCATCAaaAGTTCCAATTAAGAAGCATTTTTCA |
| B2_OvNeuroD_26_Dla1 | CTTCAGTTTCGATTAATTACTCTTTaaATCATCCAGTAAACCGCC |
| B2_OvNeuroD_26_Dla1 | CCTCGTAAATCCTCATCAaaGATAATGAAAATAATAAGCTACAAA |
| B2_OvNeuroD_26_Dla1 | CAGTTTGCTGCAATGCAATTAATTTaaATCATCCAGTAAACCGCC |
| B2_OvNeuroD_26_Dla1 | CCTCGTAAATCCTCATCAaaATGTCTCTGTTGATTTCACTTGAGT |
| B2_OvNeuroD_26_Dla1 | GAAGTTAATAAATGTGTTGCTCAGCaaATCATCCAGTAAACCGCC |
