## Supplementary General Information for "Optimization of Whole Mount RNA multiplexed in situ Hybridization Chain Reaction with Immunohistochemistry, Clearing and Imaging to visualize octopus neurogenesis"

Supplementary Material

### 1 Supplementary Figures

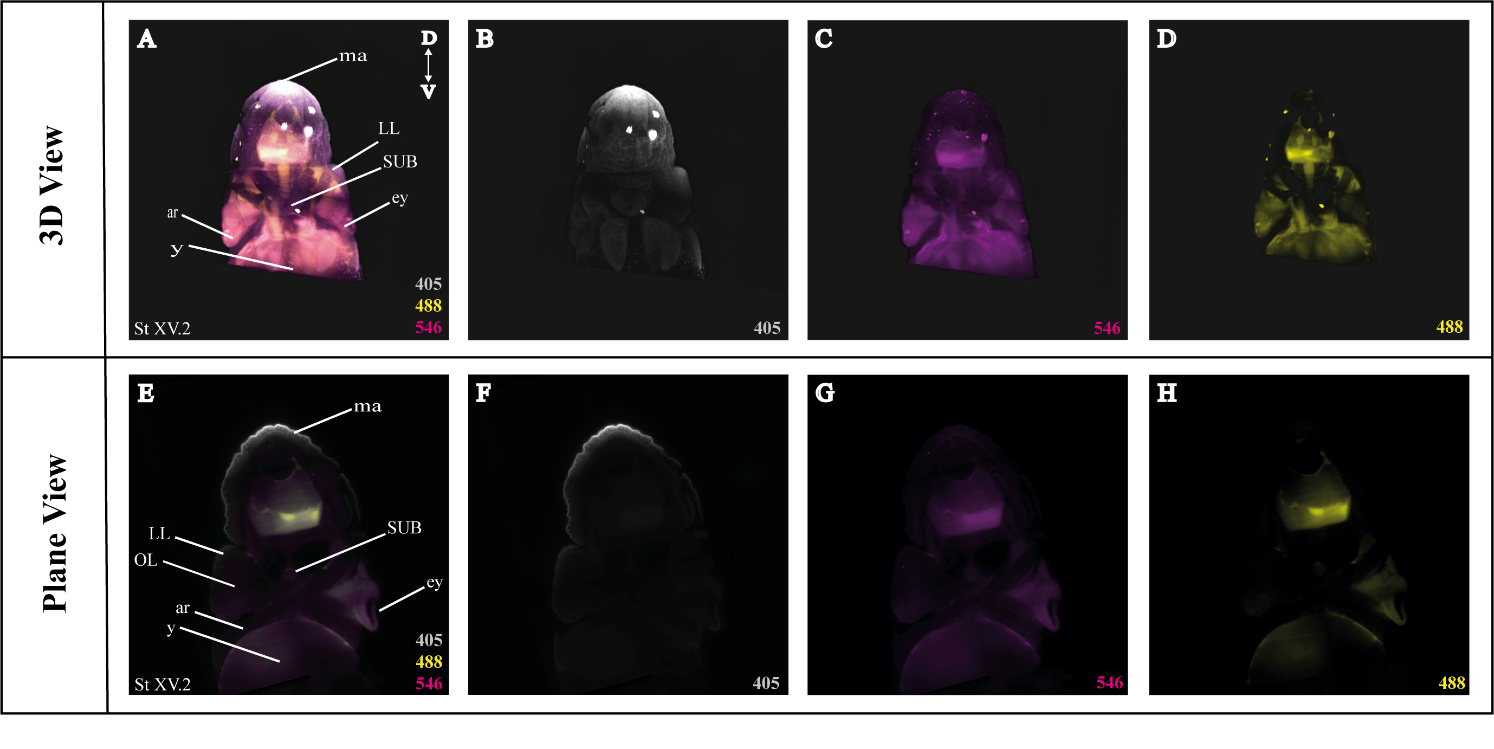

**Supplementary Figure F1.** Control Figure for Whole Mount multiplexed HCR v3.0 followed by Fructose-Glycerol Clearing on an *Octopus vulgaris* embryo (developmental stage XV) imaged with LSFM to visualize background fluorescence. Therefore, only amplifier hairpins for B1 (546) and B3 (488) were added (No probe, No DAPI). (A) Overview image showing all channels together in 3D view. (B) 405. (C) 488. (D) 546 (E) Overview image showing all the channels on a coronal section of Stage XV embryo. (F) 405. (G) 488. (H) 546 *Abbreviations: ar, arm ; D: dorsal ; ey, eye ; fu, funnel ; LL, lateral lip ; ma, mantle ; OL, optic lobe ; SUB, subesophageal mass ; V: ventral; y, yolk.*

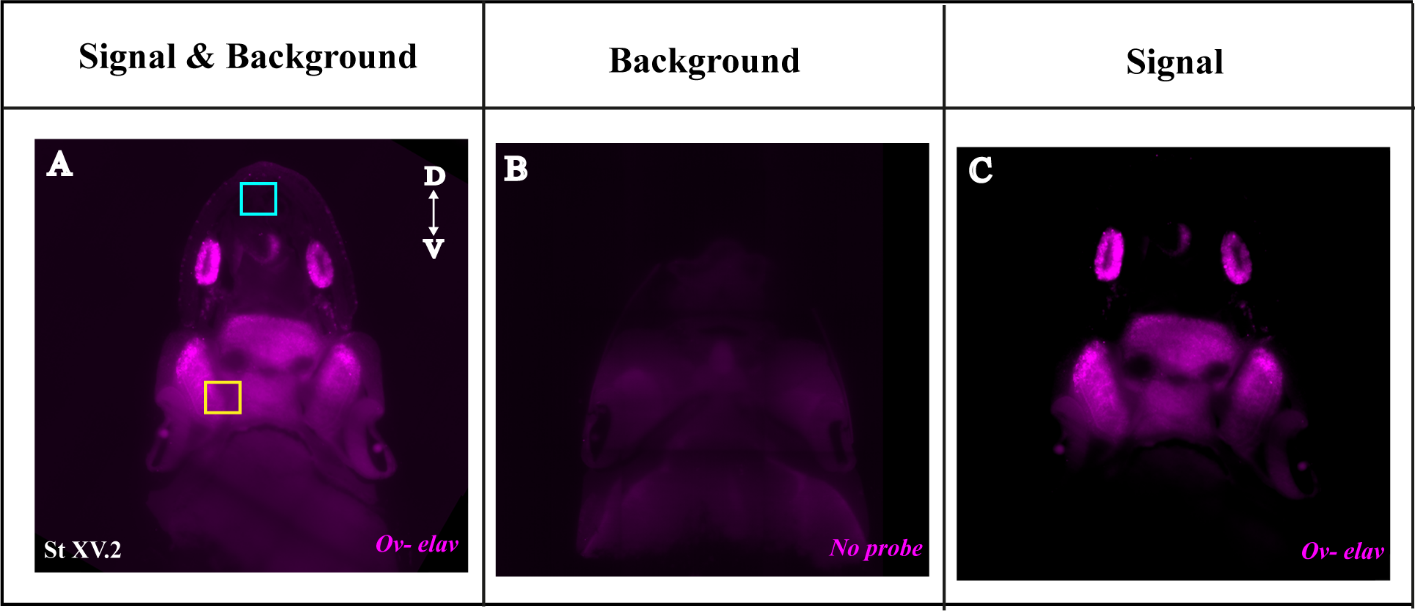

**Supplementary Figure F2.** Removing background and optimizing signal-to-background ratio of light sheet images using ARIVIS software. (A) Unedited image of *Ov-elav* on an Octopus vulgaris embryo (developmental stage XV) on a coronal section. Blue square indicates background and orange square indicates presence of signal and background. (B) Image of control embryo to which only amplifier hairpins for B1 (546) and B3 (488) were added (No probe, No DAPI). (C) Edited image showing the optimized signal-to-background ratio for *Ov-elav.*

### 2 Supplementary Videos

**Supplementary Video V1.** Video of DAPI stained stage XV octopus embryo which shows the manually reconstructed central brain and stellate ganglia. Color legend. light blue, optic lobes; pink, supraesophageal mass; orange, subesophageal mass; and dark blue, stellate ganglia.

**Supplementary Video V2.** Video of complete z-stack images of a stage XV octopus embryo in coronal plane illustrating multiplexed HCR for *Ov-elav* (magenta), *Ov-apolpp* (yellow) as well as nuclear marker DAPI (grey).

**Supplementary Video V3.** Video of complete z-stack images of a stage XV octopus embryo in coronal plane showing each channel separately illustrating multiplexed HCR for *Ov-elav* (magenta), *Ov-apolpp* (yellow) as well as nuclear marker DAPI (grey).’

**Supplementary Video V4.** Video of stage XV octopus embryo in 3D view illustrating multiplexed HCR for *Ov-elav* (magenta), *Ov-apolpp* (yellow) as well as nuclear marker DAPI (grey).

**Supplementary Video V5.** Video of complete z-stack images of a stage XV octopus embryo in coronal plane illustrating multiplexed HCR for *Ov-ascl1* (magenta), *Ov-neurod* (cyan), IHC for PH3 (yellow) as well as nuclear marker DAPI (grey).

**Supplementary Video V6.** Video of complete z-stack images of a stage XV octopus embryo in coronal plane showing each channel separately illustrating multiplexed HCR for *Ov-ascl1* (magenta), *Ov-neurod* (cyan), IHC for PH3 (yellow) as well as nuclear marker DAPI (grey).

**Supplementary Video V7.** Video of stage XV octopus embryo in 3D view illustrating multiplexed HCR for *Ov-ascl1* (magenta), *Ov-neurod* (cyan), IHC for PH3 (yellow) as well as nuclear marker DAPI (grey).

### 3 Supplementary Table

**Supplementary Table T1.** Nucleotide sequences of HCR probes
